# SpineJ : A software tool for quantitative analysis of nanoscale spine morphology

**DOI:** 10.1101/764548

**Authors:** Florian Levet, Jan Tønnesen, Valentin Nägerl, Jean-Baptiste Sibarita

## Abstract

Super-resolution microscopy provides diffraction-unlimited optical access to the intricate morphology of neurons in living brain tissue, resolving their finest structural details, which are critical for neuronal function. However, as existing image analysis software tools have been developed for diffraction-limited images, they are generally not well suited for quantifying nanoscale structures like dendritic spines. We present SpineJ, a semi-automatic ImageJ plugin that is specifically designed for this purpose. SpineJ comes with an intuitive and user-friendly graphical user interface, facilitating fast, accurate, and unbiased analysis of spine morphology.

## 1. Introduction

Dendritic spines are the postsynaptic component of most excitatory synapses in the mammalian brain. Highly recognizable in light microscopic images as protrusions covering the membrane surface of dendrites, they serve as a convenient morphological proxy for synapses (1).

Spine structure and synapse function are closely related; large spine heads contain larger postsynaptic densities (PSD) and form synapses that have a higher conductance, while activity-dependent synaptic plasticity is associated with changes in the number, size and shape of dendritic spines. Hence, spine morphology and its dynamics can give important indications about the functional state and adaptations of synapses and neuronal circuits associated with brain development, learning and memory (2; 3) and brain disorders (4).

However, dendritic spines feature nanoscale structural details, such as spine necks, which are functionally very important but cannot be properly resolved by conventional optical techniques like confocal and two-photon microscopy. Ranging in length between 0.2 and 2 micrometers with diameters smaller than 200 nm, spine necks have traditionally defied geometric analyses in live tissue.

Given this technical limitation, researchers have instead used calibrated fluorescence measurements to estimate spine size. This is in principle possible because the spatially integrated fluorescence signal scales with the volume of the source of the fluorescence, even if the underlying compartment cannot be spatially resolved (5; 6). However, the problem is that this method is indirect and requires measurements of absolute fluorescence intensity, which is sensitive to potential confounders like fluorescence bleaching, variations in laser power, and the presence of organelles or other structures in the spines (7). While electron microscopy readily provides the required spatial resolution for direct geometric measurements, its workflow is very time-consuming, subject to fixation artifacts and incompatible with live-cell analysis.

By permitting live-cell imaging with nanoscale resolution, super-resolution fluorescence microscopy is capable of providing new insights into the functional properties of spines (8). Among the new techniques increasingly adopted by neuroscientists, stimulated emission depletion (STED) microscopy is particularly well suited for imaging cellular morphology and has been successfully applied to study the structural dynamics of spines in brain slices (9; 10) and in vivo (11; 12; 13). It is a volumetric, laser-scanning imaging technique with high optical sectioning, just like confocal or 2-photon microscopy, which works with the same organic dyes and fluorescent proteins as these well-established techniques.

Several software packages are available for analyzing neuronal morphology, including dendritic spines. While these programs can determine spine densities and coarse spine size parameters, such as total length or volume, they are in general poorly suited for extracting geometric information about spine necks. A main drawback is that these programs mostly rely on data extraction after processing steps such as thresholding and binarizing (14; 15; 16; 17; 18; 19; 20; 21), which can seriously compromise the morphological information contained in super-resolution images (7).

Given the wealth of morphological data available in super-resolution microscopy images, there is a need for dedicated analysis tools that are capable of faithfully extracting the relevant nanoscale information in absolute units of physical size. Unfortunately, such tools are currently not available and manual analysis is the current working standard for the analysis of super-resolved images of spine morphology. This leaves the quantitative analysis of large data sets time-consuming and subject to user fatigue and bias. Moreover, with the manual approach researchers tend to under-utilize their data, reporting only a single neck width measurement per spine, instead of carrying out a more comprehensive analysis to capture the variation of the diameter along the neck.

## 2. Workflow

We introduce SpineJ, a sophisticated yet user-friendly ImageJ (22) plugin for semi-automatic quantification of spine morphology. SpineJ allows robust spine morphology analysis using a combination of advanced filtering and segmentation techniques accessible through a simple graphical user interface (GUI). Its workflow is composed of three main steps (**Fig. 1**) : (i) Interactive wavelet-based filtering to binarize the dendritic structures of interest from the background. (ii) Semi-automatic reconnection of spines that may have been erroneously separated from the dendritic shaft, which can happen with very thin and weakly fluorescent necks. (iii) Skeletonization-based segmentation allowing quantitative morphological analysis of spine neck width, length and head surface.

**Figure 1:**
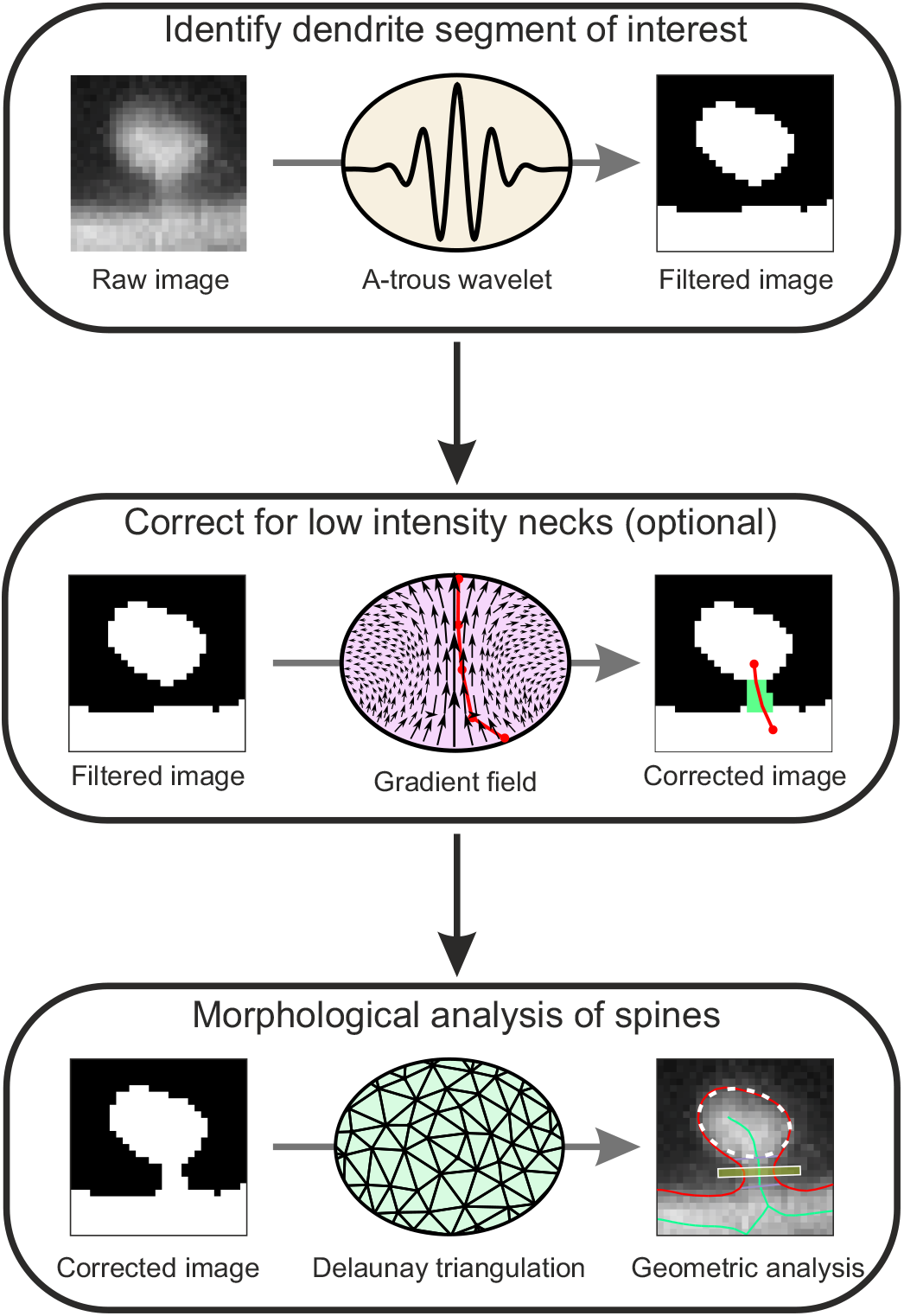
SpineJ workflow. The workflow is composed of three steps. First, the dendrites of interest are identified by using a wavelet filtering process. Spines separated from the dendrite because of weak fluorescent necks can be reconnected by following a gradient field computed on the original image. Finally, the morphological analysis of spines is performed by computing a constrained Delaunay triangulation on the dendrite outlines.

SpineJ only requires basic ImageJ skills and comes with an intuitive and user-friendly GUI that allows swift collection of spine geometry data in a reproducible and unbiased manner. The semi-automated analysis is insensitive to pixel size and shows excellent agreement with manual analyses of the same spines.

## 3. Integrated software components

### 3.1. Wavelet-based image filtering

The segmentation of spine morphology requires efficient and reliable discrimination of the fluorescent structures of interest from the background. However, in the case of dendritic spines, common segmentation techniques, such as basic thresholding or unsharp masking, are not well suited because of the differences in fluorescence intensity between the thin spine neck (100–300*nm*) and the much wider dendritic trunk (1 − 3*μm*). To overcome this problem, we used wavelet filtering, which has intrinsic multi-scale properties and allows to efficiently segment structures with various sizes and intensities.

We used the fast “à-trous” algorithm (23), which computes a series of multi-scale wavelet coefficients by iterative convolutions of increasing kernels (**Supplementary Fig. 1A, Methods**). A-trous wavelets have several key features : (i) the noise variance 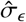 can be robustly estimated (24); (ii) the size of filtered objects is directly related to the wavelet scale, allowing segmenting structures of similar size by simply thresholding a given wavelet sub-band; (iii) wavelets are not sensitive to the absolute image intensities, making it possible to quantify and compare efficiently different images.

All the STED images analyzed here were thresholded using the second and third wavelet coefficients with a threshold of 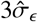 (**Supplementary Fig. 1B, Methods**). It is important to note that 2 the choice of wavelet coefficients and thresholds will mostly affect the number of spines that will have to be manually reconnected to the dendrite, but will not influence the measurements of spine neck widths, which are carried out on the raw images.

### 3.2. Spine head reconnection

Because of the low fluorescence intensity of spine necks relative to spine heads and the dendritic trunk, some spine heads may be disconnected from the parent dendrite (**Fig. 2A**). As spine necks are rarely straight and extend at variable angles from the dendrite, their reconnection using a straight line is a poor estimate of the true situation.

**Figure 2:**
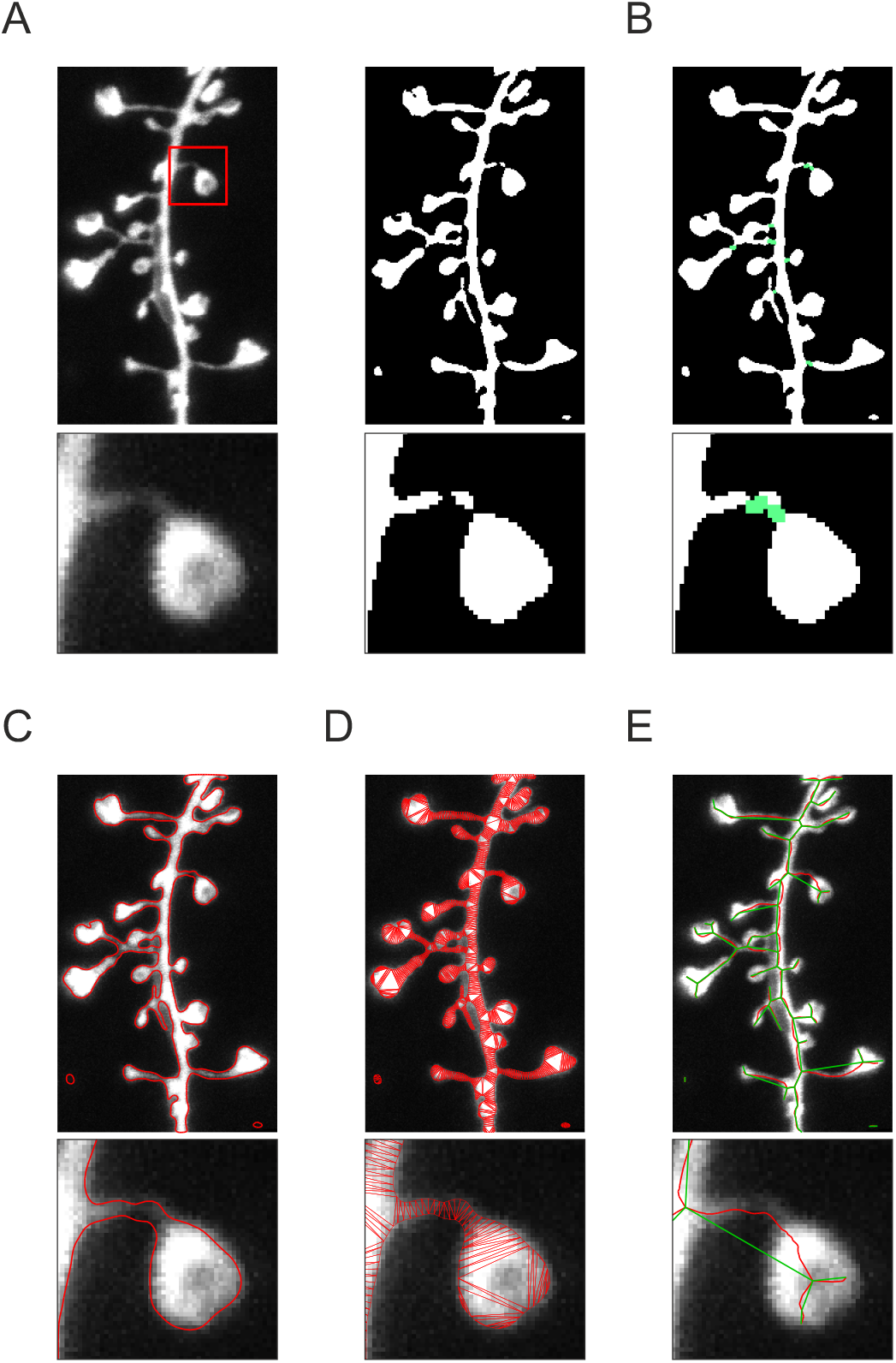
Abstracting dendrites as graphs. (A) Original STED image (left) and binarized dendrite after à-trous wavelet filtering (right). (B) Reconnection of separated dendritic spines (green). (C) Vectorial outline of the dendrite (red). (D) Constrained Delaunay triangulation computed on the vectorial outline (red). (E) Skeleton (red) and graph (green) extracted from the triangulation.

We adopted an approach originally developed in NeuronJ that allows the identification and tracing of dim structures (25). A gradient field is directly computed from the original image, resulting in pair-pixel vectors reflecting their direction and magnitude. It allows determining the best path between two points by capturing and following the orientation of bright structures present in the image. Reconnections are performed locally after manual identification of both the isolated spine heads and the parent dendrite (**Fig. 2B**).

### 3.3. Automatic spine identification

At this step, neurites are delimited from the background as a binary image and direct identification of spines is difficult as individual pixels lack context. To overcome this problem, we used skeletons, which are 1D geometric descriptors that naturally contain information on the shape of the structures.

In order to properly account for spine neck geometries, it is essential to ensure smooth vectorial skeletons. This eliminates pixel-based skeletons resulting from segmentation mask thinning since they are usually jagged. We therefore used polygon-based skeletons, where the binarized neurite is represented as a polygon. We combined the C1-continuous Catmull-Rom spline (26) with an arc-length parametrization, providing higher accuracy for curved and junctional parts of the dendrites (**Fig. 2C**), and resulting in segmenting the dendrite outline as a smooth vectorial line independent of pixel size.

The points of the spline are then used as seeds to compute a constrained Delaunay triangulation (**Fig. 2D**). This space-subdividing technique has two advantages : first, triangle connectivity can be used to extract a skeleton, with its branching and ending points defining a graph **G** that accurately describes neuronal morphology (**Fig. 2E, Methods**). We used a pruning algorithm to remove insignificant small branches from the skeleton (**Methods**). Second, a morphological compartment can be represented as a subgraph of **G** combined with a set of triangles, facilitating its geometric definition and analysis (**Fig. 3A**). In particular, we use three morphological compartments, namely dendrite, spine head, and spine neck (**Fig. 3B**). The automatic identification of spines is achieved by graph theory applied to **G** (**Methods**). In instances where spines are ill-defined, a correction can easily be manually applied using the *ROI tools* in ImageJ.

**Figure 3:**
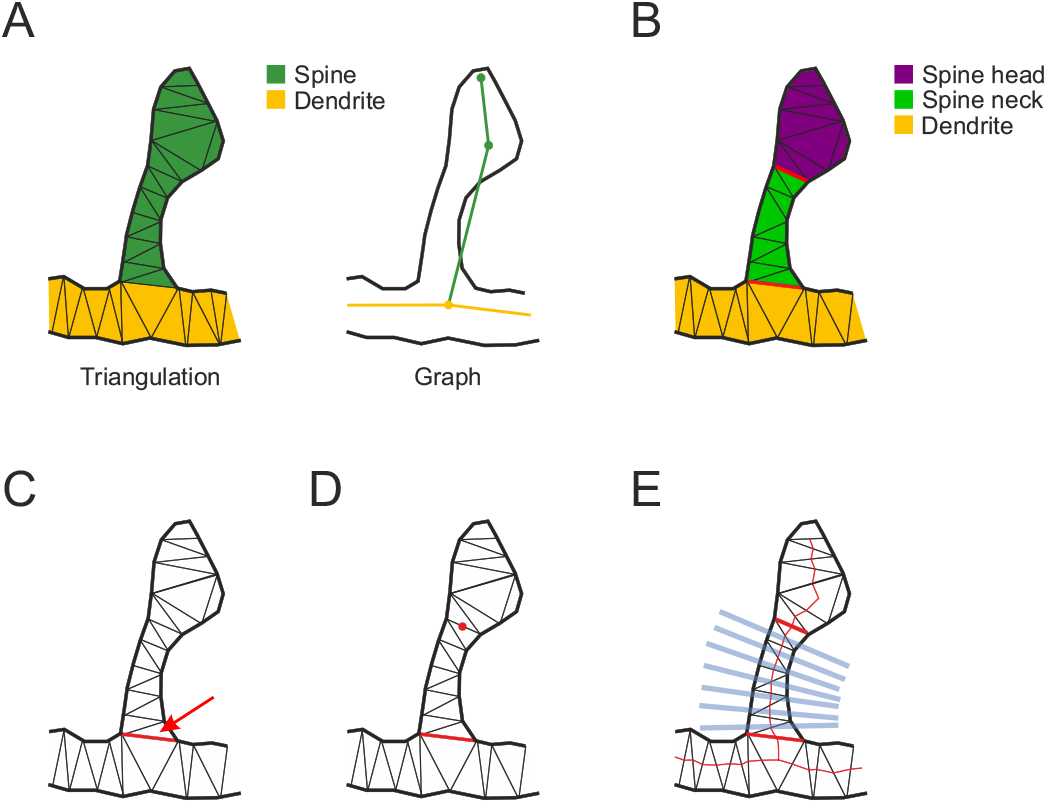
Spine neck quantification. (A) Triangle sets (left) and corresponding graph (right) of a spine (green) connected to its dendrite (orange). (B) Definition of the three morphological compartments : the spine head (magenta) and neck (green) and the dendrite (orange). (C) Basis of the spine neck is defined as the edge connecting the triangle sets of the spine and the dendrite (red arrow). (D) The user defines the neck end by clicking on a triangle edge (red circle). (D) SpineJ computes the three compartments and automatically traces several perpendicular lines to analyze the neck morphology (blue).

### 3.4. Spine quantification

Because the border between a spine head and its neck is not well defined, their separation is hard to automatize. We used the Delaunay triangulation to facilitate the separation. The neck base is defined as the shared edge between a spine and its connecting dendrite (**Fig. 3C**), while its tip, which connects the spine head, is manually defined by selecting a triangle edge (**Fig. 3D**).

The software then automatically traces evenly spaced lines perpendicular to the neck skeleton (**Fig. 3E**). Neck widths are extracted from the full width at half maximum (FWHM) computed by Gaussian fits of intensity line profiles gathered from the raw images. The spine head area is directly computed on the binarized image after separation from the neck. Most of the parameters can be selected automatically or adjusted manually (**Methods**).

## 4. Results

### 4.1. Measurements of spine neck width

Spine neck widths in STED images have usually been determined by fitting one-dimensional Gaussian (27; 28) or Lorentzian functions (29; 30) to intensity line profiles of spine neck cross sections and extracting the FWHM value. Usually, only a single measurement per spine is taken at the place where the neck appears to be the thinnest. By contrast, SpineJ automatically computes the minimal, maximal and average FWHM values all along the spine neck.

While both Gaussian and Lorentzian functions produced highly correlated measurements (*R*2 = 0.83, *y* = 0.89*x* + 9.3; **Fig. 4B**), the Gaussian fit returned slightly larger values for the neck width than the Lorentzian fit (mean_Lorentz_ 137 ± 37*nm* (SD), mean_Gauss_ 144 ± 38*nm*, p = 0.0006, paired t-test, n = 69 spines from 6 dendritic segments; **Fig. 4C**). All subsequent measurements were performed by using Gaussian fits.

**Figure 4:**
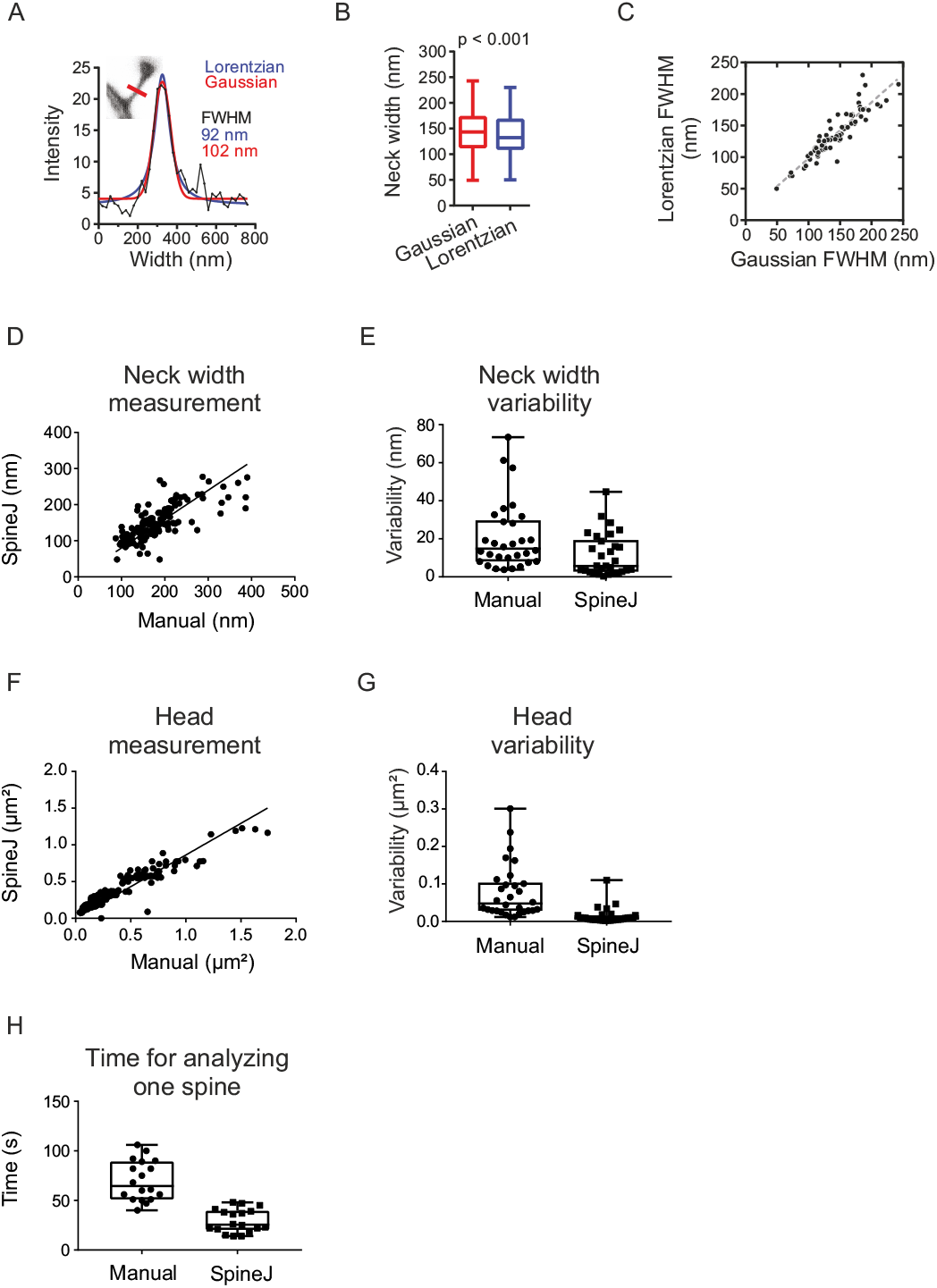
Comparison of Lorentzian and Gaussian fits and measurement of spines. (A-C) Comparison of Lorentzian and Gaussian fits. (A) Lorentzian (FWHM 92*nm*) and Gaussian (FWHM 102*nm*) fits of the same spine neck width. (B) Gaussian fit (144 ± 38*nm* SD) identifies a slightly wider width than the Lorentzian one (137 ± 37*nm* SD), n = 69 spines. (C) Correlation between the Lorentzian and the Gaussian FWHM (*R*2 = 0.83). (D-H) Comparison of manual and SpineJ analysis. (D) Correlation between the measurements obtained manually or by SpineJ for the neck width. (E) User variability of the manual and SpineJ analysis for the neck width. (F) Correlation between the measurements obtained manually or by SpineJ for the head area. (G) User variability of the manual and SpineJ analysis for the head area. (H) Comparison of the time needed to analyze one spine manually and with SpineJ. N = 150 spines.

### 4.2. Comparison of SpineJ performances with manual quantifications

We compared the performance of SpineJ with manual quantifications (**Fig. 4D-H**). We asked 5 “näıve” persons to analyze 30 spines from 3 dendritic segments. In the case of manual analysis, the angle, the position of the line across the necks and the elliptical ROI representing the spine head were set by the experimenter.

For spine neck widths, even though the values obtained with SpineJ were smaller compared to manually (SpineJ : 146 ± 4*nm* SEM; manual : 177 ± 5*nm* SEM), they were highly correlated (*y* = 0.799*x*; **Fig. 4D**). The variability between users was almost two times less with SpineJ compared to the manual analysis (SpineJ : 11 ± 2*nm* SEM; Manual : 21 ± 3*nm* SEM; **Fig. 4E**). This is expected since SpineJ computes the minimal width from many different measurements sampled all along the spine neck.

For spine heads, both SpineJ and manual analyses returned similar areas (SpineJ : 0.37 ± 0.02*μm*^2^ SEM; manual : 0.38 ± 0.03*μm*^2^), which were highly correlated (y = 0.86x; **Fig. 4F**). The variability between users was more than five times lower with SpineJ compared to manual analysis (SpineJ : 0.014 ± 0.004*μm*^2^ SEM; manual : 0.078 ± 0.013*μm*^2^ SEM; **Fig. 4G**).

Finally, the analysis time per spine was less than half for SpineJ compared to manual analysis (SpineJ : 29 ± 3*s* SEM; manual : 70 ± 5*s* SEM; **Fig. 4H**).

## 5. Discussion

By breaking the diffraction barrier, super-resolution fluorescence microscopy is providing optical access to micro-anatomical structures in live brain tissue. This has allowed geometric analysis of dendritic spines and axons, providing new insights into their biological function (10; 31). However, geometric analysis of dendritic spines in super-resolution images is currently still a manual process, which inevitably introduces variability and bias, and is very time-consuming.

Here, we introduce SpineJ, a new software to extract geometric information about nanoscale details of dendritic spines. The strength of SpineJ lies in its ability to analyze spine neck geometry in a quick and reproducible manner, returning measurements in absolute units of size, which was not possible before.

The software is based on a structured workflow design, where all user inputs are related to spine selection, after which automatic measurements are directly obtained from the original image. This sequence separation of spine selection and analyses provides a strong protection against user bias, as validated by the low variability of measurements performed by different users.

We validated the performance of SpineJ by comparing it with manual analysis. Importantly, SpineJ systematically reported a smaller minimal neck width than manual analysis, reflecting the fact that SpineJ systematically measures neck width all along the full extent of the neck and thus can reliably find the true minimal value, unlike manual analysis, which is a more “hit-or-miss” approach.

In addition, the high variability associated with manual analysis of spine heads shows that elliptical shapes are a poor representation for spine heads. Users could use the hand-drawn ROI tool to properly account for discontinuous borders, but the task would then become even more complex and lengthy. By contrast, SpineJ achieves a robust estimation of spine heads without making any assumptions about their shapes. Finally, combining robust image processing techniques with very simple user-interactions (a few mouse clicks) and instant visual feedback (preview, overlay, statistics) minimizes the time spent on analyzing each spine. While this gain in time was already significant during our tests with a rather small dataset (30 spines in 3 dendritic segments), it would surely increase with user fatigue when analyzing hundreds of spines.

Contrary to manual analysis, which is designed for a specific task and cannot be easily extended, the workflow used in SpineJ allows going beyond measurements used in the literature. The custom of measuring only a single value for neck width simply reflects a practical inadequacy of manual analysis. By contrast, SpineJ gives the complete numbers on the neck, facilitating much more quantitative and robust analyses of spine morphology (**Supplementary Fig. 2**).

We encourage researchers to adopt and use SpineJ as a reference software for more meaningful and transparent quantitative analysis of spine morphology.

## 6. Methods

### 6.1. A-trous wavelet filtering

The ‘a trous’ wavelet transform represents a discrete and translation-invariant approach to the classical continuous wavelet transform. We define *c*_0_(*k*) as the original fluorescent image. The smoothed data *c*_*i*_(*k*) at a given resolution *i* and at pixel *k* are obtained by the convolution *c*_*i*_(*k*) = ∑_*l*_ *h*(*l*)*c*_*i*−1_(*k* + 2^*i*−1^*l*), where *h* is a low-pass scaling function (usually a *B*^3^ spline). The difference between two consecutive resolution levels *w*_*i*_(*k*) = *c*_*i*−1_(*k*) − *c*_*i*_(*k*) represents the wavelet coefficients (or subband) at level *i*. Segmentation is achieved by thresholding these wavelet coefficients independently.

In fluorescence microscopy, the noise is a mixture of Gaussian (electronic) and Poisson (photons) statistics. Interestingly, during the wavelet transform, while the noise *ϵ* is decomposed in all of the wavelet sub-bands, more than 80% of its components are present in the first wavelet sub-band. As a result, (24) proposed a robust estimation of the noise varianc 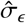 based on the median absolute value of the first wavelet coefficients and defined as 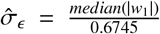. Since most of the noise *ϵ* is part of *w*_1_ while containing little useful signal, such estimator has become very popular and is widely used.

To identify pixels that are part of the actual dendrite, the wavelets sub-bands are thresholded with 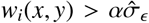 where α is a coefficient set by the user. The final binary image is reconstructed by summing all the filtered wavelet coefficient subbands. We experimentally determined that a threshold of 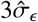 applied on the second and third wavelet sub-bands provides a robust segmentation of all our images. These values might slightly fluctuate depending on the acquisition parameters.

### 6.2. Gradient field computation and reconnection

In computer vision, ridges define a set of curves that represent local maxima of the image in at least one dimension. Since neurites are thin bright structures standing out from a darker background, they can readily be represented by ridges. Ridge detection is facilitated by computing a vector field on the image, with each pixel being associated with a vector direction and magnitude (25).

Reconnection between spine head and dendrite is initiated by two user-defined clicks within the respective structures to be reconnected. The algorithm computes the optimal path connecting the two points as the path exhibiting the minimal cumulative cost when following the vector field main directions. The ridge connects the points of maximal magnitude.

### 6.3. Polygon-based skeleton generation

Triangles of the constrained Delaunay triangulation are divided into three categories (**Supplementary Fig. 3A**) : extremal (E, one neighboring triangle), transitional (T, two neighboring triangles), and junctional (J, more than two neighboring triangles). The skeleton is obtained by connecting the midpoints of the edges shared by transitional triangles (**Supplementary Fig. 3B**). To avoid insignificant and unstable branches, a pruning algorithm is applied on the skeleton extremal branches (**Supplementary Fig. 3C**). Finally, the graph representation is easily generated by connecting all the junctional and extremal triangles, following the respective skeleton branches (**Supplementary Fig. 3D**).

### 6.4. Pruning of the skeleton

To avoid insignificant and unstable branches, extremal triangles are subject to a pruning algorithm designed to merge extremal regions with adjacent transitional triangles. If all the outline points of the extremal region are inside the semicircle, whose diameter is the edge between the extremal region and its adjacent transitional triangle, this edge is removed and the transitional triangle is added to the extremal region (**Supplementary Fig. 3C**, *top*). This process is repeated until at least one outline point is outside the semicircle or if a junctional triangle is reached (**Supplementary Fig. 3C**, *bottom*). After pruning, the skeleton is more stable and still embeds the topology of the neurite.

### 6.5. Automatic spine identification

Since the graph **G** is extracted from the Delaunay triangulation, any of its vertex or edge is linked to a set of triangles. The degree *d*(*v*_*i*_) counts the number of vertices connected to *v*_*i*_. Graph vertices are classified in 3 categories : leaf (*d*(*v*_*i*_) = 1), transitional (*d*(*v*_*i*_) = 2) or junctional (*d*(*v*_*i*_) = 3). A vertex is part of a spine if it is a leaf, or a transitional with one of its connected vertices being a leaf, or a junctional with at least two of the connected vertices being either a leaf or a transitional already identified as part of the spine (**Supplementary Fig. 4A**). By adding all the edges of **G** that connect two spine vertices plus the connected edges with only one dendrite vertex, we define a subgraph **G**_**spines**_ that represents all the spines (**Supplementary Fig. 4B**), with individual spines as the connected components of **G**_**spines**_. Each spine is then defined as the set of triangles of all its vertices and edges (**Supplementary Fig. 4C**).

### 6.6 Automatic determination of the neck width

To quantify the width along a given spine neck, SpineJ automatically traces several evenly spaced lines perpendicularly through the spine neck skeleton. Each line is defined by a width and a thickness, describing a rectangle under which SpineJ computes the intensity profile of the spine neck in the raw image, thus corresponding to line intensity profiles in previous manual analyses. In SpineJ the line thickness can be user-chosen, and here we used a fixed value of 100 nm for all our images. The number of lines *nb* is defined as 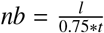, with *l* and *t* being respectively the neck length and line thickness. This ensures optimal sampling with some overlap between the lines. Each line represents the width at a specific location of the neck skeleton. This width is determined by Gaussian fitting the intensity profile with width ranging between 400 nm to 800 nm, with 50 nm steps. The fit having the highest *R*^2^ is selected as the optimal width for this skeleton position.

In our study, the final neck width reported is defined as the minimal width of all the lines computed alongside the neck skeleton.

---

**Supplementary Figure 1:**
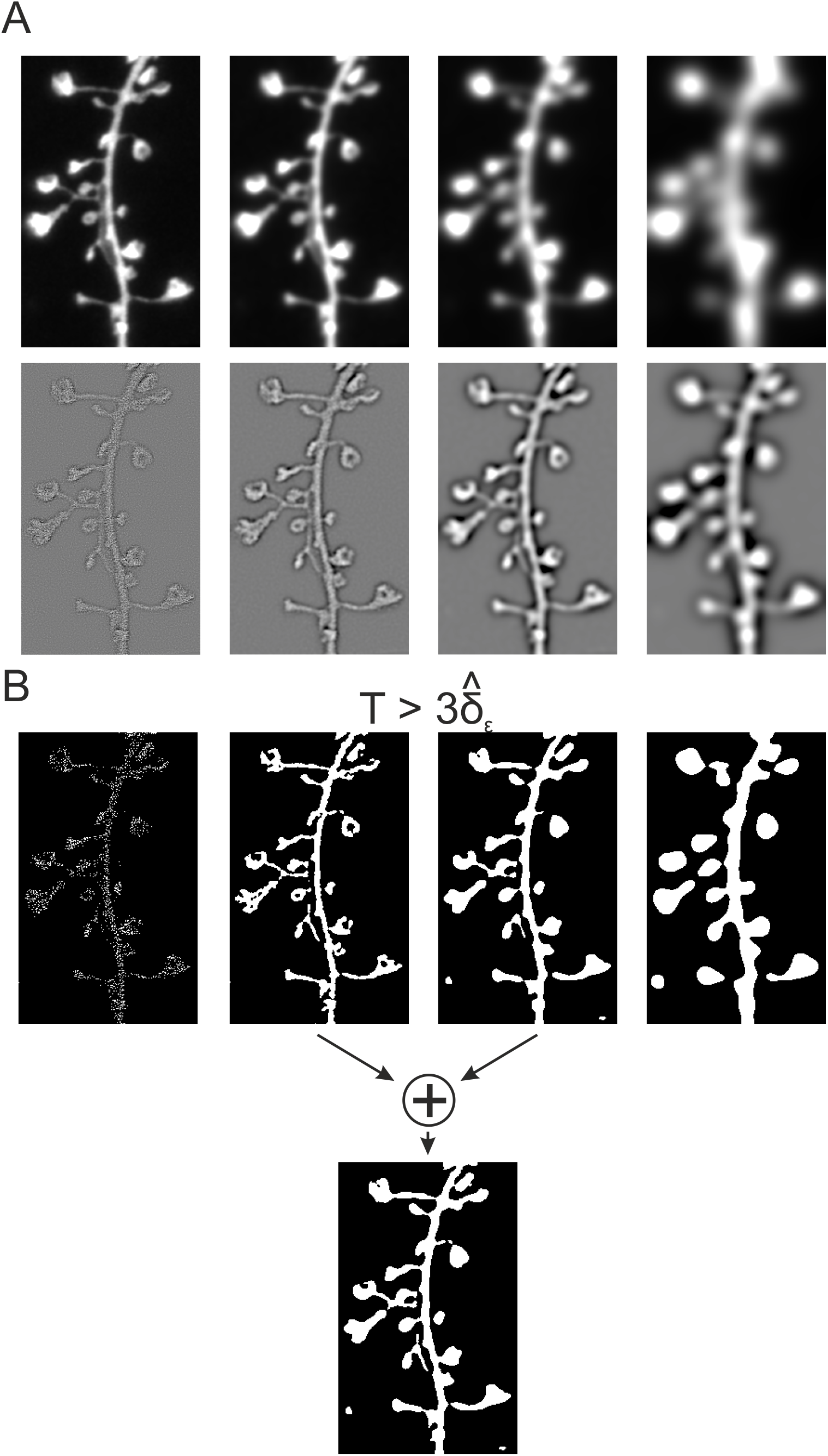
Wavelet filtering. Wavelets (A) and coefficients (B) sub-bands. (B) Filtered sub-bands resulting from applying a threshold of 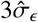 (top), with 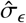 the noise variance determined automatically form the first wavelet sub-band. The final filtered image is obtained by adding the 2nd and 3rd filtered sub-bands (bottom).

**Supplementary Figure 2:**
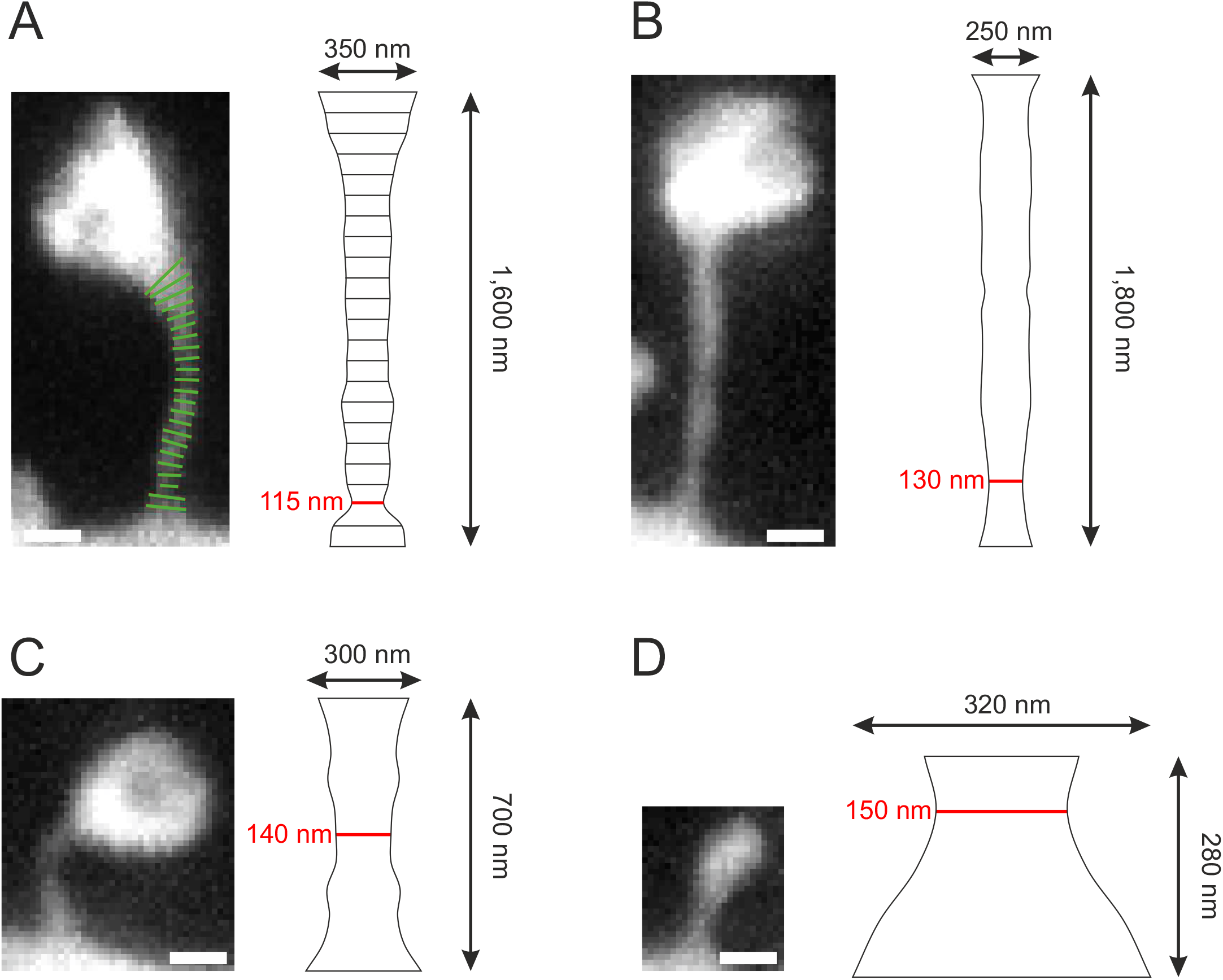
Modeling spine necks. (A) Measurements of the neck width at several locations (green) on the original image (left). (Right) Corresponding unfolding and modeling of the spine neck, with its length, maximal and minimal width (red). (B-C) modeling of the neck for 3 spines. Scale bar 400 nm.

**Supplementary Figure 3:**
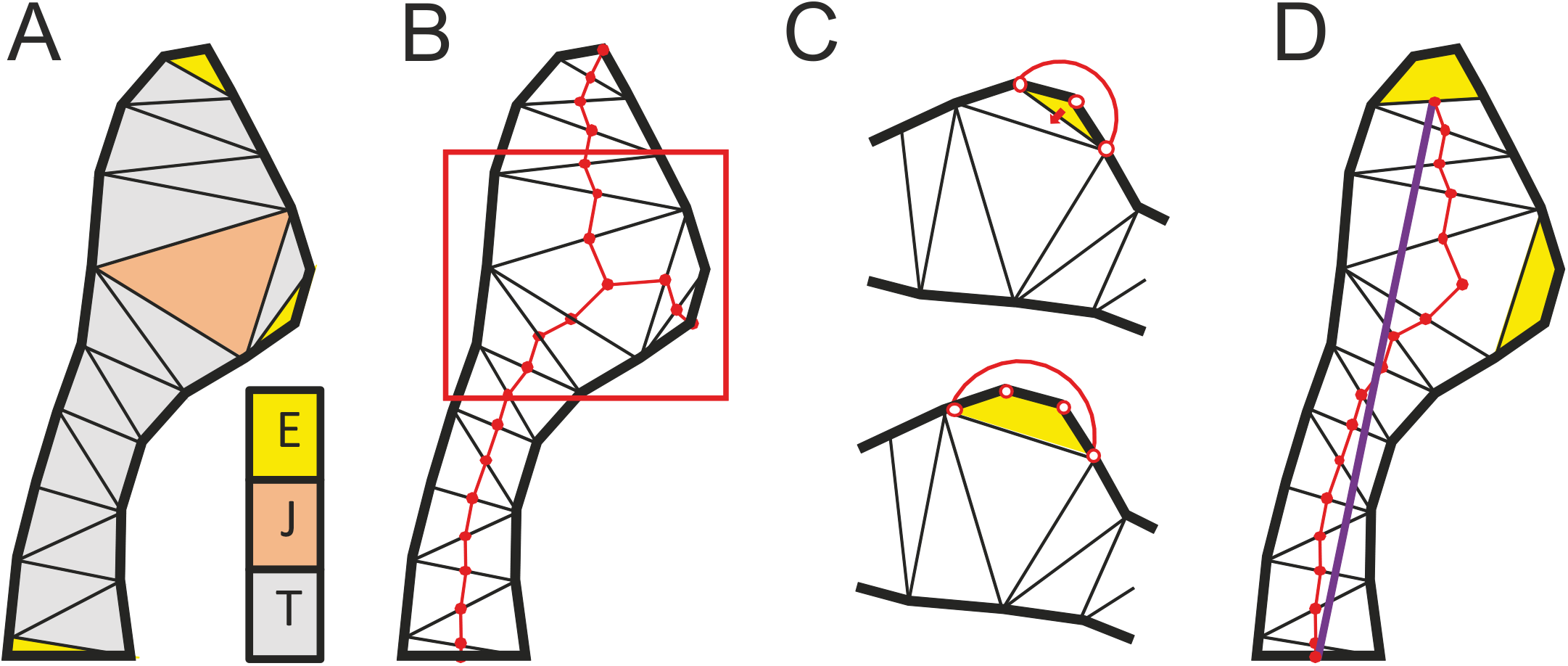
From triangulation to spine neck quantification. (A) Delaunay triangulation of a spine. Triangles are classified into three categories : extremal (E), junctional (J) and transitional (T). (B) Original skeleton extracted from the triangulation. (C) Pruning algorithm : if all points of an extremal region are inside the circumcircle, the current triangle is merged to its neighbor. This process is stopped when encountering a junctional triangle or if a point is outside the circumcircle. (D) Skeleton after pruning.

**Supplementary Figure 4:**
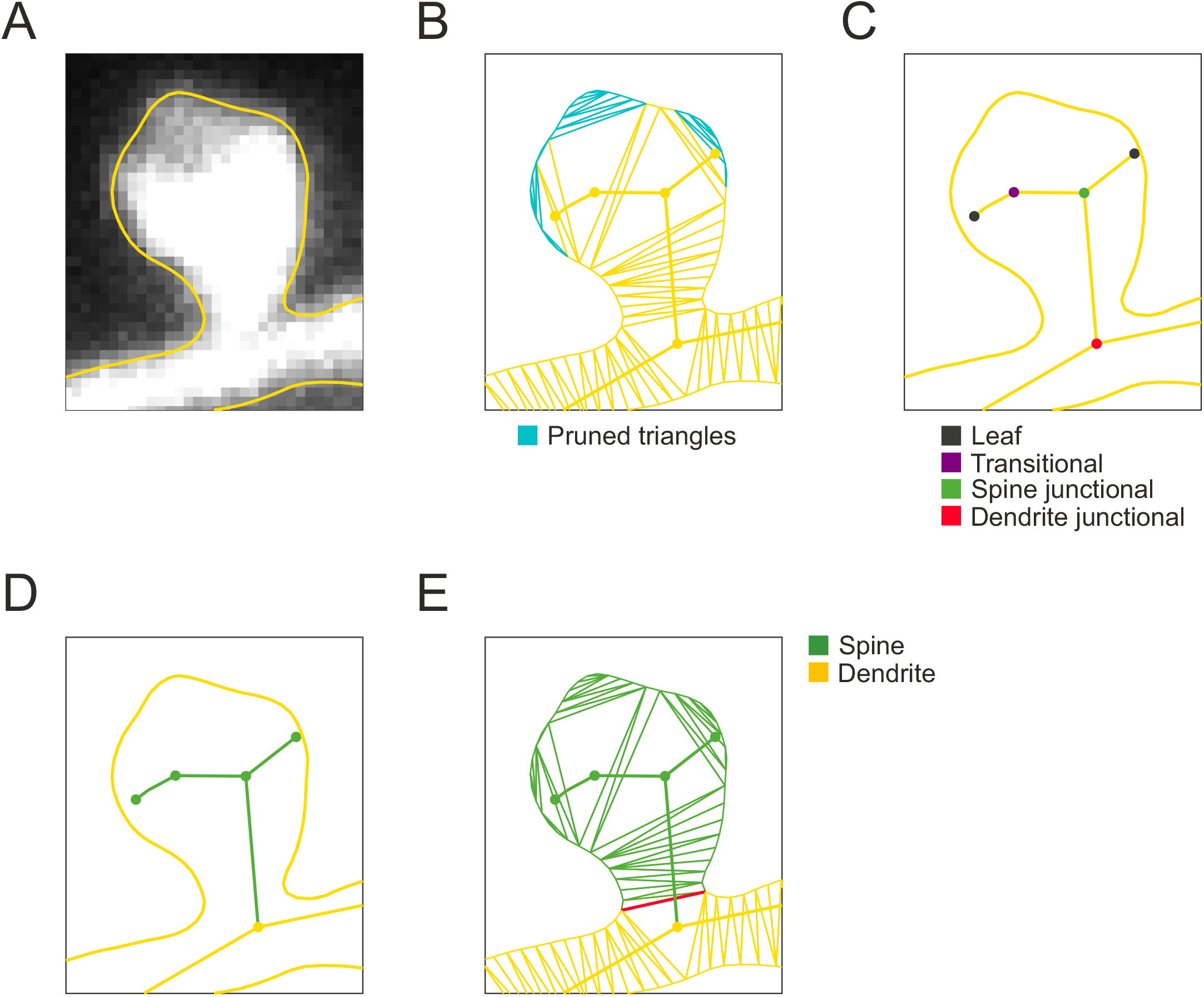
Automatic spine identification. (A) Vectorial outline of a spine and a portion of the dendrite. (B) Constrained Delaunay triangulation of the outline. The graph is computed after pruning (cyan). (C) Four classes organization of the graph vertices. (D) Subgraph representing the spine (green). It is composed of fours vertices and four edges. (E) Corresponding identified spine. The spine/dendrite separation is defined by the red edge.

## Références

[1] Sala C, Segal M (2014) Dendritic spines : The locus of structural and functional plasticity. Physiological Reviews 94(1) :141–188, DOI 10.1152/physrev.00012.2013, pMID : 24382885

[2] Yuste R, Bonhoeffer T (2004) Genesis of dendritic spines : insights from ultrastructural and imaging studies. Nature Reviews Neuroscience 5(1) :24–34, DOI 10.1038/nrn1300

[3] Holtmaat A, Svoboda K (2009) Experience-dependent structural synaptic plasticity in the mammalian brain. Nature Reviews Neuroscience 10 :647 EP –, URL https://doi.org/10.1038/nrn2699, review Article

[4] Montagna E, Dorostkar MM, Herms J (2017) The role of app in structural spine plasticity. Frontiers in Molecular Neuroscience 10 :136, DOI 10.3389/fnmol.2017.00136

[5] Svoboda K, Tank DW, Denk W (1996) Direct measurement of coupling between dendritic spines and shafts. Science 272(5262) :716–719, DOI 10.1126/science.272.5262.716

[6] Matsuzaki M, Ellis-Davies GCR, Nemoto T, Miyashita Y, Iino M, Kasai H (2001) Dendritic spine geometry is critical for ampa receptor expression in hippocampal ca1 pyramidal neurons. Nature Neuroscience 4(11) :1086–1092, DOI 10.1038/nn736

[7] Lenz MO, Tønnesen J (2019) Considerations for Imaging and Analyzing Neural Structures by STED Microscopy, Springer New York, New York, NY, pp 29–46. DOI 10.1007/978-1-4939-9077-13

[8] Tonnesen J, Nagerl UV (2013) Superresolution imaging for neuroscience. Experimental Neurology 242 :33 – 40, DOI https://doi.org/10.1016/j.expneurol.2012.10.004, URL http://www.sciencedirect.com/science/article/pii/S0014488612003871, new imaging techniques

[9] Nägerl UV, Willig KI, Hein B, Hell SW, Bonhoeffer T (2008) Live-cell imaging of dendritic spines by sted microscopy. Proceedings of the National Academy of Sciences 105(48) :18982–18987, DOI 10.1073/pnas.0810028105

[10] Tønnesen J, Katona G, Rózsa B, Nägerl UV (2014) Spine neck plasticity regulates compartmentalization of synapses. Nature Neuroscience 17 :678–685, article

[11] Berning S, Willig KI, Steffens H, Dibaj P, Hell SW (2012) Nanoscopy in a living mouse brain. Science 335(6068) :551–551, DOI 10.1126/science.1215369

[12] Willig K, Steffens H, Gregor C, Herholt A, Rossner M, Hell S (2014) Nanoscopy of filamentous actin in cortical dendrites of a living mouse. Biophysical Journal 106(1) :L01–L03, DOI 10.1016/j.bpj.2013.11.1119

[13] Pfeiffer T, Poll S, Bancelin S, Angibaud J, Inavalli VK, Keppler K, Mittag M, Fuhrmann M, Nagerl UV (2018) Chronic 2p-sted imaging reveals high turnover of dendritic spines in the hippocampus in vivo. eLife 7 :e34700, DOI 10.7554/eLife.34700

[14] Broser P, Schulte R, Roth A, Helmchen F, Waters J, Lang S, Sakmann B, Wittum G (2004) Nonlinear anisotropic diffusion filtering of three-dimensional image data from two-photon microscopy. Journal of Biomedical Optics 9(6) :1253 – 1264 – 12, DOI 10.1117/1.1806832, URL https://doi.org/10.1117/1.1806832

[15] Weaver CM, Hof PR, Wearne SL, Lindquist WB (2004) Automated algorithms for multiscale morphometry of neuronal dendrites. Neural Computation 16(7) :1353–1383, DOI 10.1162/089976604323057425

[16] Bai W, Zhou X, Ji L, Cheng J, Wong STC (2007) Automatic dendritic spine analysis in two-photon laser scanning microscopy images. Cytometry Part A 71A(10) :818–826, DOI 10.1002/cyto.a.20431

[17] Rodriguez A, Ehlenberger DB, Dickstein DL, Hof PR, Wearne SL (2008) Automated three-dimensional detection and shape classification of dendritic spines from fluorescence microscopy images. PLOS ONE 3(4) :1–12, DOI 10.1371/journal.pone.0001997, URL https://doi.org/10.1371/journal.pone.0001997

[18] Shen H, Sesack SR, Toda S, Kalivas PW (2008) Automated quantification of dendritic spine density and spine head diameter in medium spiny neurons of the nucleus accumbens. Brain Structure and Function 213(1) :149–157, DOI 10.1007/s00429-008-0184-2

[19] Janoos F, Mosaliganti K, Xu X, Machiraju R, Huang K, Wong ST (2009) Robust 3d reconstruction and identification of dendritic spines from optical microscopy imaging. Medical Image Analysis 13(1) :167 – 179, DOI https://doi.org/10.1016/j.media.2008.06.019

[20] Jungblut D, Wittum G, Vlachos A, Schuldt G, Zahn N, Deller T (2012) Spinelab : tool for three-dimensional reconstruction of neuronal cell morphology. Journal of Biomedical Optics 17(7) :1 – 8 – 8, DOI 10.1117/1.JBO.17.7.076007, URL https://doi.org/10.1117/1.JBO.17.7.076007

[21] Blumer C, Vivien C, Genoud C, Perez-Alvarez A, Wiegert JS, Vetter T, Oertner TG (2015) Automated analysis of spine dynamics on live ca1 pyramidal cells. Medical Image Analysis 19(1) :87 – 97, DOI https://doi.org/10.1016/j.media.2014.09.004, URL http://www.sciencedirect.com/science/article/pii/S1361841514001418

[22] Schneider CA, Rasband WS, Eliceiri KW (2012) Nih image to imagej : 25 years of image analysis. Nature Methods 9 :671 – 675, URL https://doi.org/10.1038/nmeth.2089

[23] Holschneider M, Kronland-Martinet R, Morlet J, Tchamitchian P (1990) A real-time algorithm for signal analysis with the help of the wavelet transform. In : Wavelets, Springer Berlin Heidelberg, pp 286–297

[24] Donoho D, Johnstone IM (1995) Adapting to unknown smoothness via wavelet shrinkage. Journal of the American Statistical Association 90 :1200–1224

[25] Meijering E, Jacob M, Sarria JC, Steiner P, Hirling H, Unser M (2004) Design and validation of a tool for neurite tracing and analysis in fluorescence microscopy images. Cytometry Part A 58A(2) :167–176, DOI 10.1002/cyto.a.20022

[26] Farin G (1993) Curves and Surfaces for Computer-Aided Geometric Design (Third Edition), third edition edn. Academic Press, DOI https://doi.org/10.1016/B978-0-12-249052-1.50005-2

[27] Ding JB, Takasaki KT, Sabatini BL (2009) Supraresolution imaging in brain slices using stimulated-emission depletion two-photon laser scanning microscopy. Neuron 63(4) :429–437, DOI 10.1016/j.neuron.2009.07.011

[28] Tønnesen J, Nadrigny F, Willig K, Wedlich-Söldner R, Nägerl UV (2011) Two-color sted microscopy of living synapses using a single laser-beam pair. Biophysical Journal 101(10) :2545–2552, DOI 10.1016/j.bpj.2011.10.011

[29] Bethge P, Chéreau R, Avignone E, Marsicano G, Nägerl UV (2013) Two-photon excitation sted microscopy in two colors in acute brain slices. Biophysical Journal 104(4) :778–785, DOI 10.1016/j.bpj.2012.12.054

[30] Takasaki KT, Ding JB, Sabatini BL (2013) Live-cell superresolution imaging by pulsed sted two-photon excitation microscopy. Biophysical Journal 104(4) :770–777, DOI 10.1016/j.bpj.2012.12.053

[31] Chéreau R, Saraceno GE, Angibaud J, Cattaert D, Nägerl UV (2017) Superresolution imaging reveals activity-dependent plasticity of axon morphology linked to changes in action potential conduction velocity. Proceedings of the National Academy of Sciences 114(6) :1401–1406, DOI 10.1073/pnas.1607541114

